# Training with audio and video games improves audiospatial performance in a “cocktail-party” task: A controlled intervention study in young adults

**DOI:** 10.1101/2020.11.17.386300

**Authors:** Jonathan Barend Schuchert, Jörg Lewald

## Abstract

Computer game playing has been suggested to be an effective training to enhance perceptual and cognitive abilities. Focusing on potential improvements in auditory selective spatial attention induced by computer gaming, we compared a passive waiting-control group with two gaming groups, playing either a first-person audio-only action game requiring spatial attention and sound localization or a platform side-scroller video game without audiospatial components, which has been shown to improve cognitive performance in previous studies. Prior to and immediately after game training for 1 month for at least 30 min per day (total training time ≥15 h), healthy young adults were tested in an audiospatial task simulating a “cocktail-party” situation with multiple speakers at different positions. The proportion of correct target localizations was significantly increased after audio and video gaming compared with the control group. However, there were no significant differences between gaming groups, with similarly strong effects of action audio game and non-action video game trainings on auditory selective spatial attention. Thus, it seems as if successful training of “cocktail-party” listening can be induced not only by modality-specific near-transfer learning within the audiospatial domain, but also by far transfer of trained cognitive skills across sensory modalities, which may enhance domain-general processes supporting selective attention.

## Introduction

Currently, video games are increasingly used as a training tool to enhance human cognitive functions (for review, see [1–3]). In particular, application of video games has been suggested to be beneficial for improving and preventing symptoms of neurodegenerative disorders, such as Alzheimer’s disease, for counteracting cognitive decline in healthy aging, and for cognitive enhancement in normal healthy people [4,5]. It is assumed that the effects of playing video games are related to processes of brain plasticity increasing volume of specific areas and connectivity between regions [6–8].

In particular, video game players have been shown to be generally better than non-players in perceiving small differences in grey scales, in processing speed, and visual attentional performance, such as optimized use of attentional resources, improved top-down and bottom-up attention, as well as superior selective visuo-spatial and peripheral attention [7,9–11]. Furthermore, positive effects of video games on shifting, updating, and dual processing as well as working memory performance have been reported (e.g. [8,12–14], for review, see [15–17]). Especially players of action video games (i.e., games with high speed, high information density, and often violence [14]), showed increased performance in numerous cognitive tasks of different difficulty levels (for review, see [18,19]). The results of cross-sectional studies with video game players, have been largely confirmed by controlled intervention studies using a repeated measures design (i.e., with testing before and after a period of gaming). In particular, these intervention studies demonstrated facilitation of attentional functions and spatial cognition after game training [9,10,20–23].

Video games have been shown to induce specific processes of structural brain plasticity that may be related to observations on the behavioral level. For example, in a controlled intervention study, using a simple platform video game, Kühn et al. [6] found significant gray matter increases in areas involved in spatial navigation, strategic planning, and working memory, namely hippocampus and dorsolateral prefrontal cortex. Similarly, a cross-sectional study by Kühn and Gallinat [24] demonstrated the amount of lifetime multi-genre video gaming to be positively associated with gray matter volumes of entorhinal, hippocampal, and occipital areas, thus suggesting adaptive neural plasticity related to navigation and visual attention. Also, a cross-sectional study by Tanaka et al. [25] reported significantly larger gray matter volume in right posterior parietal cortex of action video game experts compared with non-experts.

While positive effects of computer games on cognitive performance have been clearly established for the visual modality, the question of whether related effects also exist in the auditory modality has, to our knowledge, not been investigated so far. Also, whether a cross-modal transfer of training exists (that is, auditory improvement by video gaming or vice versa) is still an unresolved issue. Recently, a cross-sectional study by Stewart et al. [26] investigated effects of action video gaming on the participants’ performance in auditory cognitive and perceptual tasks, such as attention in listening, speech-in-noise perception, and listening in spatialized noise sentences. However, these authors failed to find any association between action video game play and auditory performance, although positive effects of video game play on a visual task were observed, as known from previous studies. Stewart et al. [26] concluded that action video game play does not result in cross-modal transfer learning, due to the absence of the players’ meaningful interaction with an acoustically relevant auditory environment during play. Thus, on the one hand, it might seem reasonable to assume that beneficial effects on auditory performance can only be induced by audio games. On the other hand, several studies have argued against this view, rather supporting conceptions of cross-modal or supra-modal learning. For example, Salminen et al. [27], using an adaptation paradigm with magnetoencephalography recording, found that neural auditory spatial selectivity was increased when participants were engaged in a visual task compared to passive listening. Also, Zhang et al. [28] reported improved frequency discrimination and auditory working memory when testing was alternated with playing *Tetris*, a visual puzzle game, in silence. Starting from the assumption that video-game playing may enhance probabilistic inference as a general learning mechanism, Green et al. [29] demonstrated improved performance of video-game players compared with non-players in both visual and auditory perceptual tasks requiring decision making based on probabilistic inference, thus suggesting transfer of video game-induced learning effects to the auditory modality.

Effects of playing audio games on human cognitive abilities have, until now, received only little attention. Audio games are games, in which all relevant information is provided acoustically. In previous studies, audio games have mainly been used to enhance navigation skills in blind persons [30,31]. Currently, recreational audio games increasingly incorporate elements of high speed, such that some of them can be described as action audio games. As increased visual attentional performance has been demonstrated in players of video action games (see above), one might assume that audio action game training may have similar effects in the auditory modality, but empirical research on this topic is missing so far. In a more general context, auditory training protocols developed for people suffering from hearing loss [32-34] or children [35] have been shown to be valid tools to improve auditory communication skills (for review, see [36]).

The present controlled intervention study started from the hypothesis that sensory training by playing an audio-only action game over a period of several weeks may significantly improve aspects of auditory selective spatial attention. For intervention, we chose the game *The Blind Swordsman* (Evil-Dog Productions, Montreal, Canada; http://www.evil-dog.com/the-blind-swordsman.html), in which the player had to identify a target sound source (an enemy) moving in a 3D virtual auditory environment among distractors and to guess its location, distance, and speed. To asses selective auditory spatial attention before and after the period of gaming, we used a paradigm simulating a so-called “cocktail-party” situation [37], in which participants had to localize the position of a non-verbal predefined target source among three distractor sources [38]. Given the open question of whether video games can have positive effects on auditory performance, as mentioned above, we also included a second intervention group, which played a well-established video non-action platform game *(Mega Mario;* A. Weber, Berlin, Germany; http://mmario.sourceforge.net/). Several previous studies have demonstrated non-auditory cognitive effects for this type of video game, such as enhancements of processing speed, reasoning, visuospatial coordination functions, visuomotor coordination, working memory, antisaccadic inhibition, and general cognitive performance, as well as structural brain plasticity in hippocampus, frontal eye field, and dorsolateral prefrontal cortex [6,39–42]. We hypothesized that auditory game training may result in significant enhancement of auditory selective spatial attention compared with a non-playing control group. Furthermore, if auditory performance could be exclusively improved by modality-specific training, we assumed that audio-game training will result in stronger improvements in the auditory task than video-game training over the same period. Alternatively, similar effects of both these interventions would rather argue in favor of cross-modal transfer of learning processes induced by video gaming to the auditory modality.

## Materials and Methods

A pre-post parallel-groups design was employed. Three groups of subjects were tested: (*1*) an audio-only game group; (*2*) a video game group; and (*3*) a passive control group. All subjects were tested in two experimental sessions, immediately before and after a period of about 1 month, during which both gaming groups played the assigned game daily for 30 min and the control group did not play computer games. Thus, for the two active groups the minimum amount of total game training time was 15 h.

### Subjects

Fifty-seven subjects (39 women, 18 men; mean age 23.6, SE 0.5, range 19-34 years) participated in the experiment. As assessed by the Edinburgh Handedness Inventory [43], 41 subjects were right-handed (laterality quotient, *LQ* ≥ 30), 12 were left-handed (*LQ* ≤ 30), and four were ambidextrous (|*LQ*| < 30). Subjects were randomly assigned to three groups (audio game; video game; passive control) of equal sizes, with equal proportions of women and men in each group (see below), as sex is known to be a factor in cocktail-party listening performance [44]. There were no significant differences between groups in mean age (*H(2)* = 1.72, *p* = 0.42) and mean handedness *LQ (H(2)* = 0.10,*p* = 0.95; Kruskal-Wallis tests). None of the subjects had any experience with playing audio-only games, while 49 subjects (86.0%) had experience with playing video games. Experienced gamers, playing more than 1 h per week over the last 2 months, were not recruited. All subjects had normal hearing (mean hearing level ≤ 25 dB; 0.125-8 kHz), as assessed by audiometric testing, and did not report any neurological or psychiatric disorders. In addition, a minimum of at least 70% correct responses in the single-source localization task of the first experimental session was chosen as inclusion criterion, since localization errors, such as front-back reversals and lack of externalization, can occur due to the presentation of virtual sound sources via headphones [45], as was used in the auditory tasks (see below). Twenty-four further participants were excluded from the data analysis. Of these, 13 subjects did not met the inclusion criterion of at least 70% correct responses in the single-source localization task, and eleven subjects did not complete the training. Subjects were paid for participation in the tests or received course credits. All subjects gave their written informed consent to participate in this study, which was approved by the Ethical Committee of the Faculty of Psychology of the Ruhr University Bochum. This study conformed to the Code of Ethics of the World Medical Association (Declaration of Helsinki), printed in the British Medical Journal (18 July 1964).

### Auditory tasks

During the auditory tasks, the subject sat on a comfortable chair in front of a desk in a dimly illuminated, sound-attenuating room. Auditory stimuli were presented via open, circumaural stereo headphones (HD650, Sennheiser, Wedemark, Germany). Two tasks had to be completed: (*1*) the single-source task, in which an isolated sound source had to be localized; (*2*) the multiple-sources task, in which four different sound sources were presented simultaneously at different locations, one of these a predefined target that had to be localized. The multiple-sources task largely resembled that described in [38], with the exception that we used virtual sound sources and not loudspeakers. Four different animal vocalizations (‘birds chirping’; ‘dog barking’; ‘frog’; ‘sheep’; taken from [46]), were adjusted to four different durations (300 ms; 600 ms; 900 ms; 1200 ms) by cutting out parts of the sound file using the software Cool Edit 2000 (Syntrillium Software Corporation, Phoenix, AZ, USA). For presentation of virtual sources, sound files were convolved with generic head related transfer function (HRTF) filters [47]. Using a procedure described elsewhere in detail [48,49], each sound was passed through HRTF filters delivered by Tucker-Davis Technologies (TDT, Alachua, FL, USA), using the RPvds graphical design tool software in combination with a TDT RP2.1 real-time processor system. HRTF filter coefficients were derived from measurements conducted by Gardner and Martin [50,51] with a Knowles Electronic Mannequin for Acoustic Research (KEMAR; size 14 cm from ear to ear) under anechoic conditions, and each HRTF was stored as a 256-tap FIR filter.

Virtual source locations were implemented at four different azimuth positions at 0° elevation: 60° and 20° to the left, and 20° and 60° to the right (Fig 1). Simultaneous presentation of four virtual locations in the multiple-sources task was created by digitally mixing four different waveforms, each at a different virtual location. Multiple sound sources were always presented with the same duration. Sources were either presented with identical levels for all four sound sources, or with the target source presented at a 6 dB higher or lower level with reference to the level of each of the three distractor sources. That is, target level and stimulus duration were varied between trials. This was done to have a wider range of task difficulty levels (increasing with decreasing duration and target level), as individual differences in baseline performance were relatively large (see Fig 2). Stimuli were converted to analog form via a PC-controlled, 16-bit soundcard (Audigy 2NX, Creative Labs, Singapore) and were presented at a mean sound pressure level of 62 dB(A).

**Fig 1.**
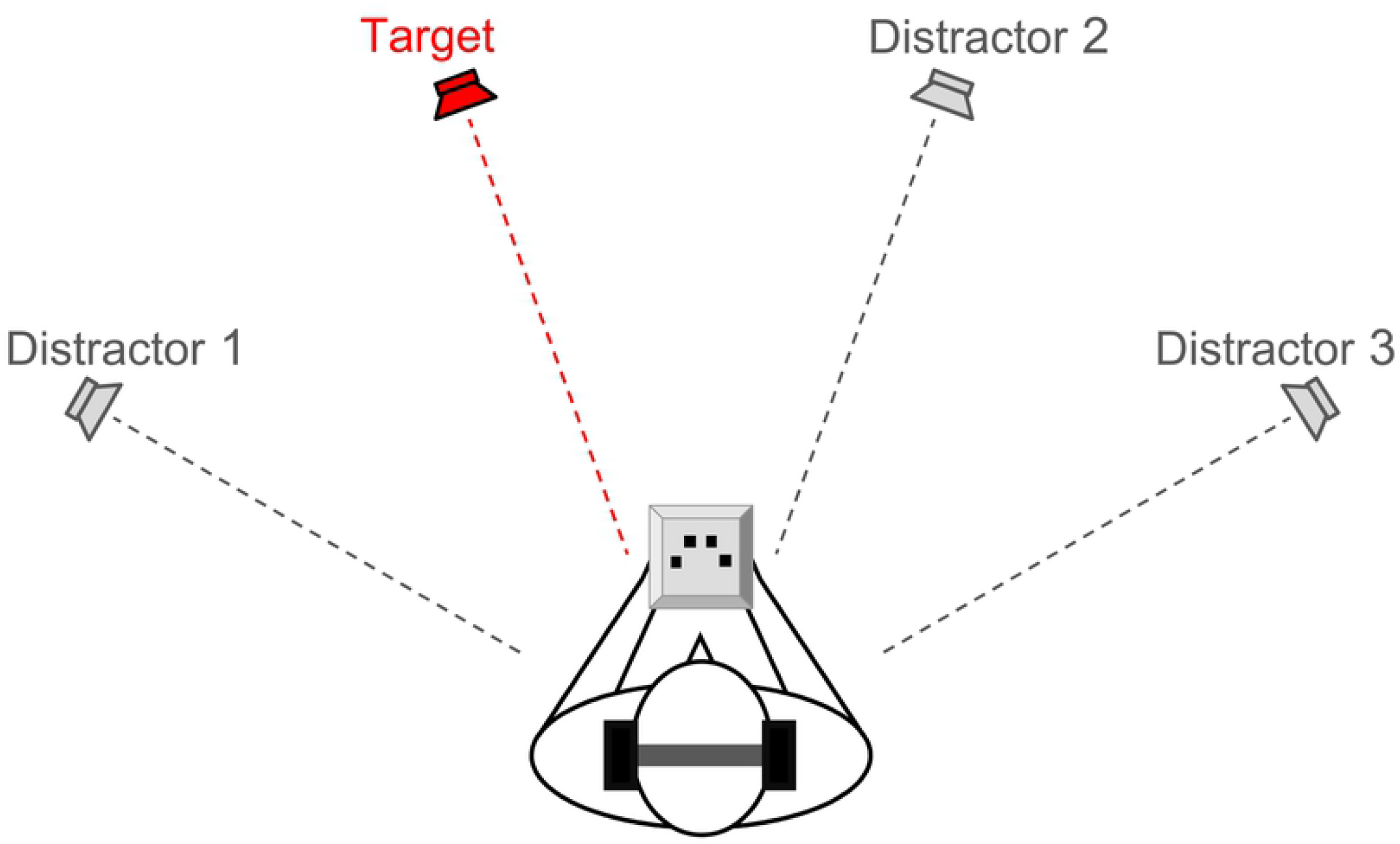
Sound-localization tasks. In the multiple-sources task, four stimuli (four different animal vocalizations, one target and three distractors) were presented simultaneously from four virtual sound locations at 60° and 20° to the left and right. Subjects were instructed to indicate the location of the predefined target vocalizations using a response box with four keys. In the single-source task, the target was presented without distractors. Apart from that both tasks were identical.

**Fig 2.**
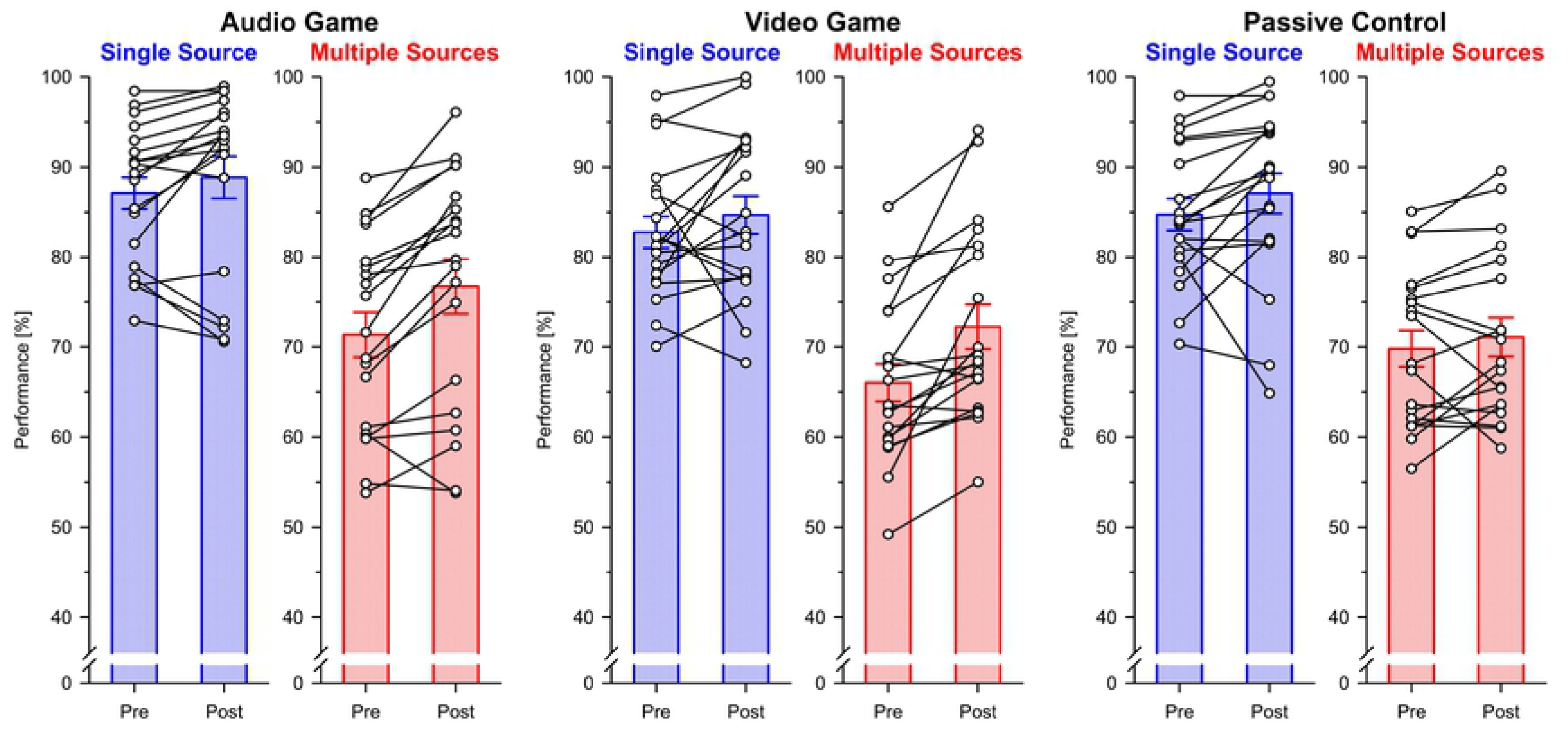
Effects of audio gaming and video gaming on sound-localization performance in single-source and multiple-sources conditions. Percentages of correct responses obtained in first (pre) and second (post) testing sessions are shown separately for the two tasks and the three groups (audio game; video game; passive control). Symbols and lines indicate individual results; bars indicate mean values (error bars, standard errors).

As in previous studies (e.g. [38]), localization performance was assessed using a spatial four-alternative forced-choice method. The subjects were informed that there were four possible positions of the target (slightly to the left; farther to the left; slightly to the right; farther to the right). They were instructed to indicate the location of the target by pressing one out of four response keys on a response box within about 1 s after each stimulus presentation. On the response box, the four keys were arranged in a semicircle, corresponding to the four possible positions of the target. Subjects were instructed to respond in each trial, without omissions, and were encouraged to guess when they were unsure about the correct position. Trial durations depended on the subject’s response time: The next stimulus was always presented 1 s after the response, thus usually resulting in trial durations of about 3 s. If the subject’s response was given earlier than 0.2 s or later than 5 s after stimulus onset, the trial was automatically repeated at the end of sub-blocks of 48 trials or at the end of the block, until the complete set of responses was recorded.

Each session comprised eight blocks, with each of the four animal vocalizations presented as target in both tasks. Each target was first presented in the single-source task and then in the multiple-sources task (with a short break between the two blocks), such that the subject was sufficiently familiar with the target when the multiple-sources task began. After the completion of the two blocks with the same target, the subject was allowed to rest for a few minutes, if required. The sequence of the targets was balanced across subjects for each group. In the single-source task, 96 trials (4 target positions *×* 3 target levels *×* 4 durations *×* 2 repetitions) were presented for each target. In the multiple-sources task, 288 trials (4 target positions × 3 target levels × 4 durations × 6 distractor combinations) were presented. In both tasks, target location, target level, and stimulus duration were varied following a fixed pseudorandom order. The timing of the auditory stimuli and the recording of the subjects’ responses were controlled by custom-written software. No feedback was given to the subjects about their performance.

### Computer Games

The audio-game group (*n* = 19; mean age 22.8 yrs, SE 0.8, range 19-30 yrs; 6 women, 13 men) was instructed to play the audio-only action game *The Blind Swordsman* (Evil-Dog Productions, Montreal, Canada; http://www.evil-dog.com/the-blind-swordsman.html). All information necessary to play this game is presented auditorily. The participant is playing from a first-person perspective as a blind swordsman who, on his quest to regain his eyesight, has to defeat enemies at several levels of increasing difficulty. The player is only able to identify the enemies’ actions and directions via auditory cues and has to react without receiving any visual information. Participants were familiarized with the game after completion of the tests in the first experimental session and received a detailed description of the game. They installed the free game software on their private PCs or laptops. The subjects were instructed to play the game daily for at least 30 min.

The audio-game group (*n* = 19; mean age 24.2 yrs, SE 0.8, range 19-30 yrs; 6 women, 13 men) was instructed to play the non-action video game *Mega Mario* (A. Weber, Berlin, Germany; http://mmario.sourceforge.net/), which is a clone of the well-known *Super Mario Bros 1* game (Nintendo, Kyoto, Japan). *Mega Mario* is a two-dimensional platform game, in which the player advances through different levels of increasing difficulty, overcoming obstacles and enemies in order to complete the main quest. The subjects were instructed to play the game daily for at least 30 min.

The passive control group (*n* = 19; mean age 23.7 yrs, SE 0.9, range 19-34 yrs; 6 women, 13 men) underwent the same procedures of testing as the gaming groups about one month apart. Control subjects did neither receive any information about the actual background of the study, nor about the fact that they were part of a control group. The actual durations of the time intervals between pre- and post-testing did not significantly differ between groups (audio group: mean 30.9 days, SE 0.6, range 28-37 days; video group: mean 30.42 days, SE 0.3, range 27-33 days; control group: mean: 31.6 days, SE 0.6, range 27-37 days; *H*(2) = 1.55, *p* = 0.46; Kruskal-Wallis test).

### Data Analysis

As in a related study [52], for the main analysis the percentages of correct responses were transformed into rationalized arcsine units (RAUs; [53,54]). RAUs have a greater range than the corresponding percent-correct scores for extreme values (< 20%; > 80%), such that the variance of RAU values is more uniform than that of percent-correct scores. Individual RAUs obtained with post-testing were normalized with reference to pre-testing. Pre-normalized RAU values were pooled across target stimuli, positions and levels and submitted to a two-factor repeated-measures ANOVA with task (single source; multiple sources) as within-subject factor and group (audio game; video game; passive control) as between-subjects factor. Post-hoc ANOVAs and *t*-tests were applied to investigate effects in detail. One-tailed testing was used for post-hoc comparisons between groups, as the primary goal was to determine if the pre-normalized performance of the audio-game group was improved compared with the video-game group and the control group. If appropriate, Bonferroni-corrected *α*-levels were used to determine statistical significance.

## Results

The participants’ percentages of correct responses assessed in the baseline sessions were analyzed using a two-factor ANOVA, with task (single source; multiple sources) as within-subject factor and group (audio game; video game; passive control) as between-subjects factor. The ANOVA did neither indicate differences between groups (*F*(2,54) = 1.69, *p* = 0.19, *η*P^2^ = 0.06), nor an interaction (*F*(2,54) = 0.41, *p* = 0.66, *η*p^2^ = 0.02). As was to be expected from the substantial differences in task difficulty, subjects performed better in the single-source, than in the multiple-sources, task (*F*(1,54) = 388.13,*p* < 0.0001, *η*p^2^ = 0.88). Individual levels of baseline performance were quite variable (Fig 2), but clearly above chance level (25%) for all subjects in both the single-source task (mean 84.86%, SE 1.03%, range 70.05–98.44%; *p* < 0.0001, binomial test) and the multiple-sources task (mean 69.05%, SE 1.29%, range 49.22–88.80%; *p* < 0.0001, binomial test).

For the main analysis, the percentages of correct responses were transformed into RAU values (see Data analysis). Then, individual data were normalized with reference to baseline performance (Fig 3). Across groups, these values were significantly above zero in both tasks, thus indicating generally better performance in the post-sessions with reference to baseline (single-source task: *t*(56) = 3.60,*p* = 0.0006; multiple-sources task: *t*(56) = 5.71,*p* < 0.0001; one-sample *t*-tests). The pre-normalized RAU values were analyzed using a two-factor repeated-measures ANOVA with task (single source; multiple sources) as within-subject factor and group (audio game; video game; passive control) as between-subjects factor. There was a significant task × group interaction (*F*(2,54) = 4.56, *p* = 0.015, *η*p^2^ = 0.14), but no main effects of task (*F*(1,54) = 1.88, *p* = 0.18, *η*p^2^ = 0.03) or group (*F*(2,54) = 0.87, *p* = 0.42, *η*_P_^2^ = 0.03). Post-hoc testing was conducted using two one-factor ANOVAs (separately for each task) with group as between-subjects factor. An effect of group was found for the multiple-sources task (*F*(2,54) = 4.47, *p* = 0.016, *η*p^2^ = 0.14; Fig 3B), but not for the single-source task (*F*(2,54) = 0.03, *p* = 0.97, *η*p^2^ < 0.01; Bonferroni-corrected *α* = 0.025; Fig 3A). Subsequent post-hoc comparisons between groups for the multiple-sources task using *t-* tests (one-tailed) revealed that both the audio-game group (mean difference 4.93 RAU, SE 1.76 RAU; *t*(36) = 2.79, *p* = 0.004, *d* = 0.91, achieved power 1 – *β* = 0.72, calculated with G*Power 3.1.9.2 [55]) and the video-game group (mean difference 5.28 RAU, SE 1.99 RAU; *t*(36) = 2.65, *p* = 0.006, *d* = 0.86, 1 – *β* = 0.67) showed stronger improvements in performance than the passive control group, while there was no significant difference between active groups (mean difference 0.35 RAU, SE 2.15 RAU; *t*(36) = 0.16, *p* = 0.44, *d* = 0.05, 1 - *β* = 0.02; Bonferroni-corrected *α* = 0.0167; Fig 3).

**Fig 3.**
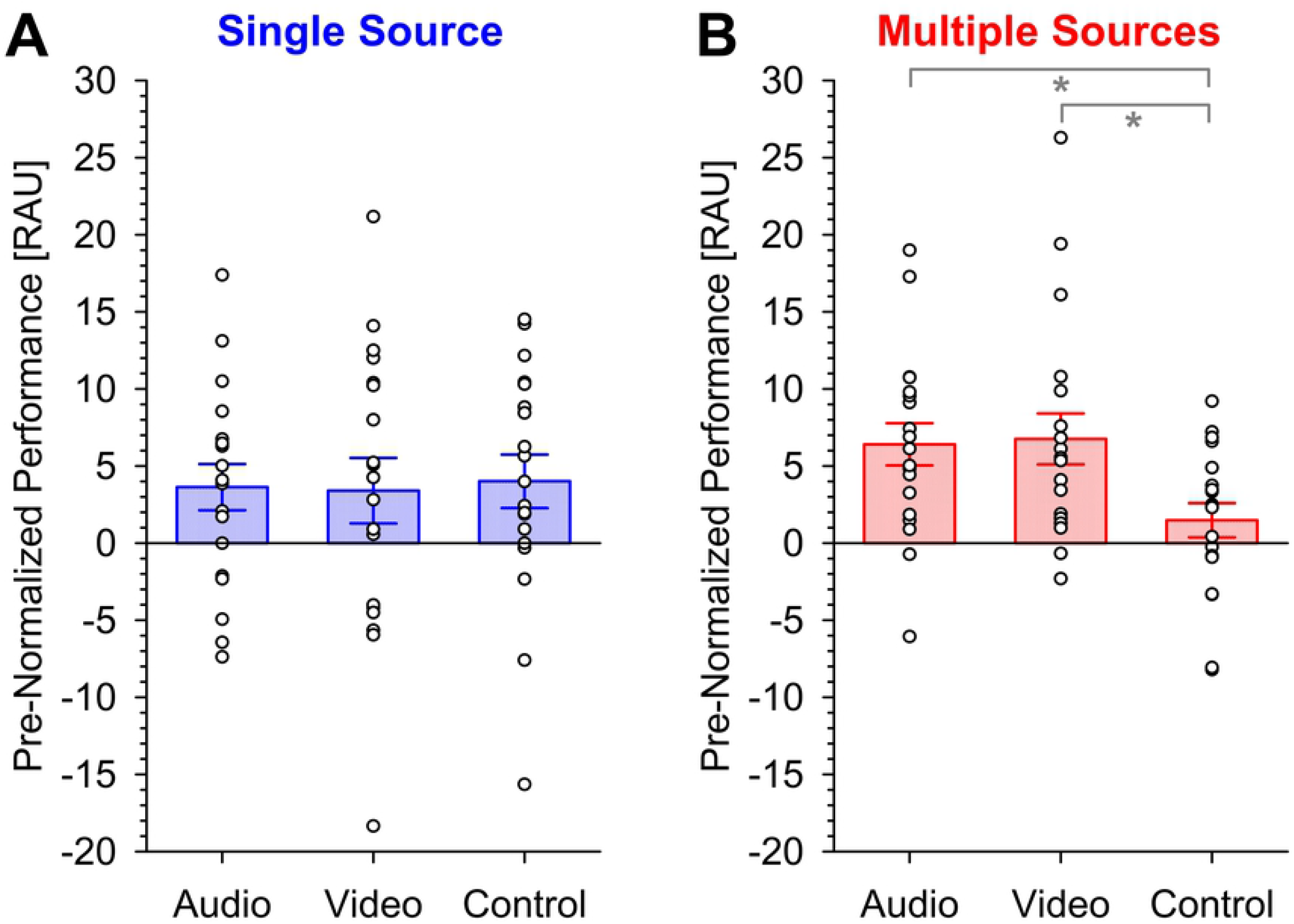
Pre-normalized performances in localization after audio and video gaming and for the passive control group. (A) Single-source condition. (B) Multiple-sources condition. The original percentages of correct responses were transformed into rationalized arcsine units (RAU values). Symbols indicate individual results; bars indicate mean values for each group (error bars, standard errors). Asterisks indicate significant improvement compared with the control group (*p* ≤ 0.006, one-tailed; Bonferroni-corrected *α* = 0.0167).

## Discussion

We found improving effects with similarly strong effect sizes of both action audio game and non-action video game training on auditory selective spatial attention, while no effects were revealed for single-source localization. There was no significant difference in improvements in multiple-sources localization obtained after both types of game training. On the one hand, these results clearly confirmed our hypothesis that playing an action audio game with spatial interaction is an effective near-transfer training enhancing audiospatial performance in complex listening situations. On the other hand, the finding that playing a non-action platform video game, which demanded spatial attention to a much lesser degree and in a different sensory modality, was about equally effective as the action audio game, was in apparent contrast to the view that learning processes during game play are modality-specific, without transfer from the visual to the auditory domain [26]. It seems as if successful training of “cocktail-party” listening depended on the enhancement of domain-general cognitive aspects of selective attention, which may have been induced by both games to a similar extent, rather than modality-specific factors of auditory spatial perception.

As a main finding, this study demonstrated for the first time a beneficial effect of playing an audio-only action game on audiospatial performance. This may parallel previous results from the visual domain, showing improving effects of action and non-action video games on visual attentional functions, in particular visual selective spatial attention [2,10,20]. Action audio game and non-action video game trainings had quite similar effects on selective auditory spatial attention. There was merely a non-significant numerical trend of stronger increase in performance after audio-game, compared with video-game, training, rather suggesting equality of effect sizes (cf. Fig 3B). This negative finding was not necessarily expected, given the recent cross-sectional study by Stewart et al. [26], who did not find any association between action video game play and auditory performance in tasks requiring attention in listening, speech-in-noise perception, and listening in spatialized noise sentences. The conclusion of these authors that action video game play does not result in cross-modal transfer learning seems to be in opposition to the result of the present intervention study, which demonstrated a causal relation of non-action video-game playing and improvement in selective auditory spatial attention. Thus, this outcome might argue in favor of a cross-modal transfer of the attentional skills trained by video gaming to the auditory domain. For the type of platform video game used here, previous research showed enhancements in several cognitive domains, such as processing speed, reasoning, visuospatial coordination functions, visuomotor coordination and working memory [39] as well as antisaccadic inhibition as a measure of frontal inhibition due to increased grey matter volume in frontal eye field [42] and increased short term memory performance and Montreal Cognitive Assessment scores in conjunction with increased hippocampal and cerebellar grey matter volumes [40] for its 3D-counterpart. In particular, there is evidence that playing a related platform video game can induce structural brain plasticity, with increases of hippocampus, entorhinal-cortex, and occipital-cortex volume found after a few months to several years of training [6,24,40,41]. Whether audio game training over longer periods can induce similar plastic changes is an open question that has to be answered empirically. The brain regions involved in the audiospatial task used here to assess selective auditory spatial attention have recently been described in great detail. The main areas were planum temporale, posterior superior temporal gyrus, inferior parietal lobule, superior parietal lobule/precuneus, inferior frontal gyrus, and dorso-frontal cortex [38,44,49,52,56–58]. Interestingly, the occipital cortex, which was shown to be increased in volume after video game playing [24], has been shown to be involved also in audiospatial functions (e.g., [59,60]). Most importantly, there is broad evidence that the frontal eye-field region is specifically concerned with functions of auditory selective attention, including audiospatial processing in “cocktail-party situations”, as was tested here [49,56,61–65]. Since the grey matter volume of frontal eye field has been found to be increased due to video game training with *Super Mario* [42] and *Tetris* [66], it seems possible that the improvement in audiospatial performance, as was observed in both the video-game and the audio-game groups of the present study, was related to plastic changes in this region. Further studies might use brain imaging techniques to investigate potential effects of audio game and video game trainings on audiospatial processing in cortical areas concerned with hearing in “cocktail-party” situations.

It has been proposed that action video games generally enhance a learning mechanism of probabilistic inference, thus allowing also for far transfer of learned cognitive skills across sensory modalities [29]. Green et al. [29] provided support for this hypothesis by demonstrating improved performance of video-game players compared with non-players in both visual and auditory perceptual tasks requiring decision making based on probabilistic inference. In this context, it has to be noted that the “cocktail-party” task used here required a spatial decision about the position of the target source presented among distractors. Thus, one could assume that improvement of probabilistic inference should have a beneficial effect on the performance in this task. Also, the two games used here may require probabilistic inference and may induce related learning processes. This may hold true not only for the action audio game, but also for the platform video game since playing requires, in either case, quick decisions in response to unforeseen events. In this regard, the present results can not only be explained by assuming that game training enhanced domain-general attentional skills related to the task, as discussed above, but also by improvement of probabilistic inference with game training. On the basis of the results, it is not possible to decide which of these explanations is more likely.

It is notable that training-induced improvements were found in the multiple-sources, but not in the single-source, condition. One possible explanation might be that game training had effects on higher-order cognitive functions of spatial hearing, as were relevant in a “cocktail-party” situation, rather than the more basic mechanisms of sound localization required for successfully completing the single-source task. The single-source task could be resolved primarily by evaluation of interaural differences in time and level and allocation of these cues to the egocentric spatial frame of reference (for review, see [45]), whereas the “cocktail-party” task was much more demanding insofar as it additionally involved processes of selective attention, in particular extraction of relevant information and inhibition of distractors. It seems as if repetitive gaming selectively modulated the latter, higher-level processes. However, one has also to consider that the performances already measured in the baseline session of the single-source task were substantially higher than in the multiple-sources task, with individual percentages of correct responses of more than 80% in the majority of participants (cf. Fig 2). Although a RAU transformation was used to correct scores for extreme values (cf. Fig 3), this null result must thus be interpreted with some caution since one cannot completely exclude that it was due to a ceiling effect.

In conclusion, we provided first evidence from data obtained in a controlled intervention study that an action audio game enhanced audiospatial performance in healthy young adults. Thus, on the one hand, action audio games may be suitable training interventions in the auditory domain. On the other hand, effects of non-action video game training were quite similar, suggesting cross-or supramodal processes associated with computer game training. The results left open the question of whether any form of computer game-based training can improve selective auditory spatial attention, independent of the sensory modality within which skills are trained, or improvements can be optimized by interventions requiring intramodal (near) transfer of learned skills. This issue has to be investigated in subsequent studies, using training interventions over longer periods than in the present study. From an application-oriented point of view, both audio and video game-based could lead to effective intervention programs for persons suffering from deficits in “cocktail-party” listening, namely healthy older people and individuals using hearing aids or cochlea implants. Also, these results suggested that action audio-only game training could be a promising tool for improving spatial abilities in blind or visually impaired persons who were unable to benefit from visual game training (cf. [30]). Moreover, it seems reasonable to assume that patients with visual field defects, such as hemianopia (cf. [67–69]), or visuospatial attention deficits, such as neglect (cf. [70]), could specifically benefit from audio-game training. This possibility should be considered by future research.

## Acknowledgements

The authors are grateful to Peter Dillmann for preparing software and stimuli used for testing. We acknowledge support by the German Federal Ministry of Education and Research in the framework of the TRAIN-STIM project (01GQ1424E) and by a donation from Sorg Hörsysteme Hörgeräte - Akustik GmbH, Schonach im Schwarzwald, Germany.

